# Apelin signaling dependent endocardial protrusions promote cardiac trabeculation in zebrafish

**DOI:** 10.1101/2021.08.30.458182

**Authors:** Jialing Qi, Annegret Rittershaus, Rashmi Priya, Shivani Mansingh, Didier Y.R. Stainier, Christian S.M. Helker

**Affiliations:** Department of Developmental Genetics, Max Planck Institute for Heart and Lung Research, 61231 Bad Nauheim, Germany; RP: The Francis Crick Institute, Organ Morphodynamics Laboratory, London NW1 1AT, UK; CSMH: Philipps-University Marburg, Faculty of Biology, Cell Signaling and Dynamics, 35043 Marburg, Germany

## Abstract

During cardiac development, endocardial cells (EdCs) produce growth factors to promote myocardial morphogenesis and growth. In particular, EdCs produce Neuregulin which is required for ventricular cardiomyocytes (CMs) to seed the multicellular ridges known as trabeculae. Defects in Neuregulin signaling, or in endocardial sprouting towards CMs, cause hypotrabeculation. However, the mechanisms underlying endocardial sprouting remain largely unknown. Here, we first show by live imaging in zebrafish embryos that EdCs interact with CMs via dynamic membrane protrusions. After touching CMs, these protrusions remain in close contact with their target despite the vigorous cardiac contractions. Loss of the CM-derived peptide Apelin, or of the Apelin receptor, which is expressed in EdCs, leads to reduced endocardial sprouting and hypotrabeculation. Mechanistically, Neuregulin signaling requires endocardial protrusions to activate extracellular signal-regulated kinase (Erk) signaling in CMs and trigger their delamination. Altogether, these data show that Apelin signaling dependent endocardial protrusions modulate CM behavior during trabeculation.

## Introduction

To meet the needs of the growing embryo, the vertebrate heart has to undergo a series of complex morphogenetic events to transform from a linear tube into a mature organ. During trabeculation, CMs in the outer curvature of the ventricles delaminate towards the lumen to form multicellular sponge-like projections, called cardiac trabeculae (Sedmera and Thomas, 1996; Sedmera et al., 2000; Stankunas et al., 2008; Liu et al., 2010; Peshkovsky et al., 2011; Staudt et al., 2014). Cardiac trabeculae are crucial to achieve increased contractility as well as for the formation of the conduction system. Trabeculation defects are often associated with left ventricular noncompaction (Oechslin et al., 2000; Claudia and Josef, 2004), embryonic heart failure, and lethality (Gassmann et al., 1995; Lee et al., 1995; Lai et al., 2010; Liu et al., 2010; Rasouli and Stainier, 2017).

In zebrafish, as in other vertebrates, the early embryonic heart consists of two monolayers of cells, the myocardium and the endocardium, that are separated by a layer of extracellular matrix (ECM) termed the cardiac jelly (CJ) (Stainier and Fishman, 1992; Brutsaert et al., 1996). Recently, it has been shown that EdCs, similar to blood endothelial cells (ECs), form sprouts, which are mostly oriented towards the myocardium (Del Monte-Nieto et al., 2018). During sprouting angiogenesis, ECs first extend filopodia to sense the microenvironment for growth factors, then they migrate into avascular areas and form new blood vessels (Gerhardt et al., 2003). Due to its similarity to sprouting angiogenesis, the sprouting of EdCs has been termed endocardial sprouting. However, whether endocardial sprouting is regulated by the same signaling pathways as sprouting angiogenesis is not known.

Multiple signaling pathways have been implicated in cardiac trabeculation, including neuregulin (Nrg)/ErbB signaling. Mouse and zebrafish embryos lacking the endocardium derived ligand Nrg or the ErbB receptor, which is expressed by the myocardium, fail to form trabeculae (Gassmann et al., 1995; Lee et al., 1995; Meyer and Birchmeier, 1995; Lai et al., 2010; Liu et al., 2010; Rasouli and Stainier, 2017). Furthermore, endocardial Notch signaling (Grego-Bessa et al., 2007; D’Amato et al., 2016; Del Monte-Nieto et al., 2018), angiopoietin 1/Tie2 signaling (Suri et al., 1996; Tachibana et al., 2005; Qu et al., 2019), and semaphorin 3E (Sema3E)/plexinD1 signaling (Sandireddy et al., 2019) are required for cardiac trabeculation in mouse. Of note, genetic deletion of the relevant receptors in the endocardium results in attenuated endocardial sprouting (Qu et al., 2019) and trabeculation defects (Grego-Bessa et al., 2007; D’Amato et al., 2016; Del Monte-Nieto et al., 2018; Qu et al., 2019; Sandireddy et al., 2019).

Cells communicate by a variety of mechanisms including paracrine and contact dependent signaling. More recently, a novel mechanism of cell communication by active transport of signaling molecules through filopodia-like actin rich membrane protrusions, also known as cytonemes, has been shown in different models including *Drosophila* (Ramirez-Weber and Kornberg, 1999; Roy et al., 2011; Huang et al., 2019), chick (Sanders et al., 2013), zebrafish (Stanganello et al., 2015), and mouse (Fierro-Gonzalez et al., 2013). Like filopodia, cytonemes depend on actin polymerization by various effector proteins including formins, profilin and IRSp53, a substrate for the insulin receptor (Rottner et al., 2017).

In this study, we take advantage of the zebrafish model, as its transparency allows single-cell resolution and high-speed imaging of the beating heart, to analyze endocardial-myocardial communication during embryogenesis. By investigating *apelin* (*apln*) mutants, we found that endocardial protrusion formation is controlled by Apln signaling. We also observed by *in vivo* imaging that endocardial protrusions promote cardiac trabeculation by modulating Nrg/ErbB/Erk signaling. Altogether, our results provide new insights into the role of endocardial protrusion during cardiac trabeculation.

## Results

### Endocardial-myocardial interactions in zebrafish

The early embryonic heart is composed of two cell types: endocardial cells and myocardial cells; and in zebrafish, myocardial cells initially form a monolayer (Figure 1A-D). In order to analyze possible interactions between the endocardial and myocardial cells, we genetically labeled the actin cytoskeleton of the endocardium using the *TgBAC(cdh5:Gal4ff)* and *Tg(UAS:LIFEACT-GFP)* lines, and the membrane of cardiomyocytes with mCherry using the *Tg(myl7:mCherry-CAAX)* line. We observed endocardial protrusions extending towards the myocardium at 24 (Figure 1A-A’’) and 48 (Figure 1B-B’’, Figure 1-figure supplement 1A) hpf. Of note, we observed more endocardial protrusions in the ventricle than in the atrium at 48, 60 and 72 hpf (Figure 1-figure supplement 1B). Subsequently, these ventricular endocardial protrusions formed anchor points with the myocardium which according to similar observations in mouse (Del Monte-Nieto et al., 2018) we refer to as touchdowns (Figure 1B-B’’). Notably, these touchdowns are stable even during cardiac contractions (Figure 1E-H, Figure 1-video 1). Starting at around 60 hpf, CMs delaminate from the compact layer towards the lumen to seed the trabecular layer (Figure 1C, C’, and C’’’’), as reported before (Liu et al., 2010; Staudt et al., 2014; Priya et al., 2020). We observed that endocardial protrusions appear to extend along the delaminating CMs (Figure 1C’’ and C’’’, Figure 1-video 2). Next, trabecular CMs start to assemble into trabecular units, which consist of several trabecular CMs, starting at 72 hpf (Figure 1D, D’, and D’’’’). At this time point, we noticed that endocardial protrusions can be detected in close proximity to trabecular CMs (Figure 1D’’ and D’’’, Figure 1-video 3). Recently, it has been shown that endothelial protrusions modulate neurogenesis by affecting progenitor proliferation in the developing brain (Di Marco et al., 2020). To determine whether endocardial protrusions affect CM proliferation, we performed EdU labeling between 28 and 72 hpf and analyzed the heart at 72 hpf. We observed that 54% of EdU positive CMs (total n=24) were in close proximity to endocardial protrusions (Figure 1-figure supplement 2) indicating that endocardial protrusions may modulate CM proliferation.

**Figure 1.**
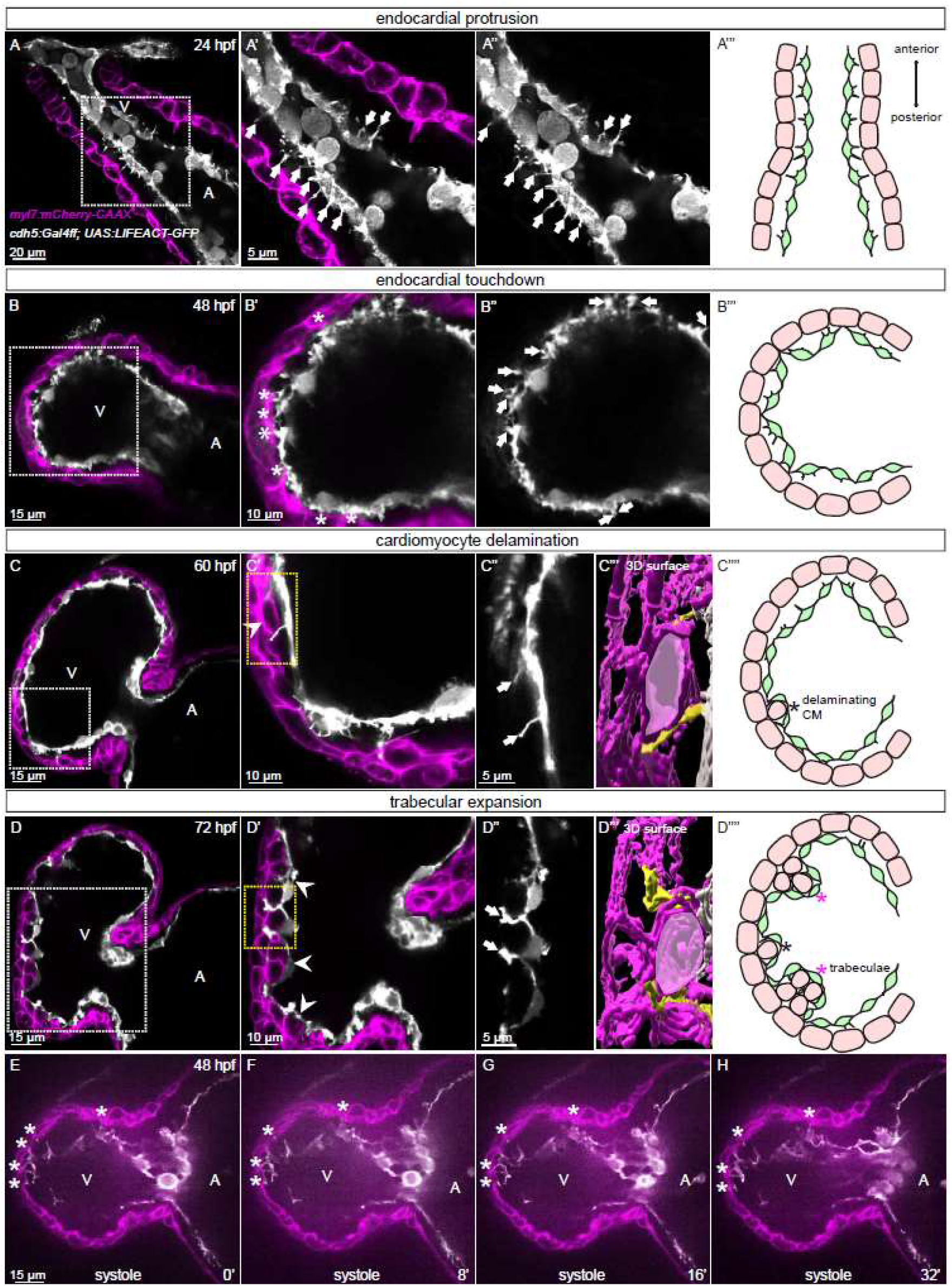
Endocardial-myocardial interactions during zebrafish heart development. **(A-D)** Confocal projection images of the heart of *Tg(myl7:mCherry-CAAX)*; *Tg(cdh5:Gal4ff)*; *Tg(UAS:LIFEACT-GFP)* zebrafish at 24 **(A)**, 48 **(B)**, 60 **(C)** and 72 **(D)** hpf. **(A-A’’)** Endocardial protrusions (arrows) towards the myocardium at 24 hpf. **(B-B’’)** Endocardial protrusions (arrows) and touchdowns (asterisks) with the myocardium at 48 hpf. **(C-C’’’)** Endocardial protrusions (arrows) during CM delamination (arrowheads) at 60 hpf. **(C’’’)** 3D surface rendering of the area in the yellow box in **C’**. **(D-D’’’)** Endocardial protrusions (arrows) during trabecular assembly and expansion (arrowheads) at 72 hpf. **(D’’’)** 3D surface rendering of the area in the yellow box in **D’**. **(A’’’’-D’’’’)** Schematics of endocardial protrusion, endocardial touchdown, CM delamination, and trabecular expansion. Black asterisks indicate delaminating CMs; purple asterisks indicate trabeculae. **(E-H)** Still images from a spinning disc time-lapse movie of a 48 hpf *Tg(myl7:mCherry-CAAX)*; *Tg(cdh5:Gal4ff)*; *Tg(UAS:LIFEACT-GFP)* heart. White asterisks indicate endocardial touchdowns. All images are ventral views, anterior to the top. V, ventricle; A, atrium.

### Genetically blocking endocardial protrusion formation reduces myocardial trabeculation

Since we observed a correlation between endocardial protrusions and myocardial trabeculation, we next aimed to examine the function of endocardial protrusions during cardiac morphogenesis. To this aim, we generated a transgenic line, *Tg(UAS: irsp53^dn^-p2a-RFP)*, to specifically block protrusion formation in the endothelium. IRSp53 regulates the actin cytoskeleton to enable cells to form different types of membrane extensions (Nakagawa et al., 2003; Millard et al., 2005; Scita et al., 2008). By crossing the *Tg(UAS: irsp53^dn^-p2a-RFP)* line to the *TgBAC(cdh5:Gal4ff)* line to overexpress Irsp53^dn^ specifically in endothelial cells, we observed a 70% reduction in the number of endocardial protrusions at 48 hpf (Figure 2A, B, and E) while their distribution appeared mostly unaffected (Figure 2A, B, and F). To test the hypothesis that endocardial protrusions modulate myocardial trabeculation, we analyzed embryos overexpressing *irsp53^dn^* in their endothelial cells in the *Tg(myl7:BFP-CAAX)*, a CM membrane line. Upon *irsp53^dn^* overexpression in ECs, we detected fewer endocardial touchdowns (Figure 2A and B). In addition, cardiac trabeculation was reduced (Figure 2C, D, G, and G’). In order to analyze a possible effect of endocardial protrusions on CM proliferation, we overexpressed *irsp53^dn^* in the endothelium in the context of the *Tg(myl7:mVenus-gmnn)* reporter to visualize cycling CMs. Compared with controls, endothelial overexpression of *irsp53^dn^* led to significantly fewer mVenus-Gmnn^+^ CMs in the ventricle (Figure 2H and I).

**Figure 2.**
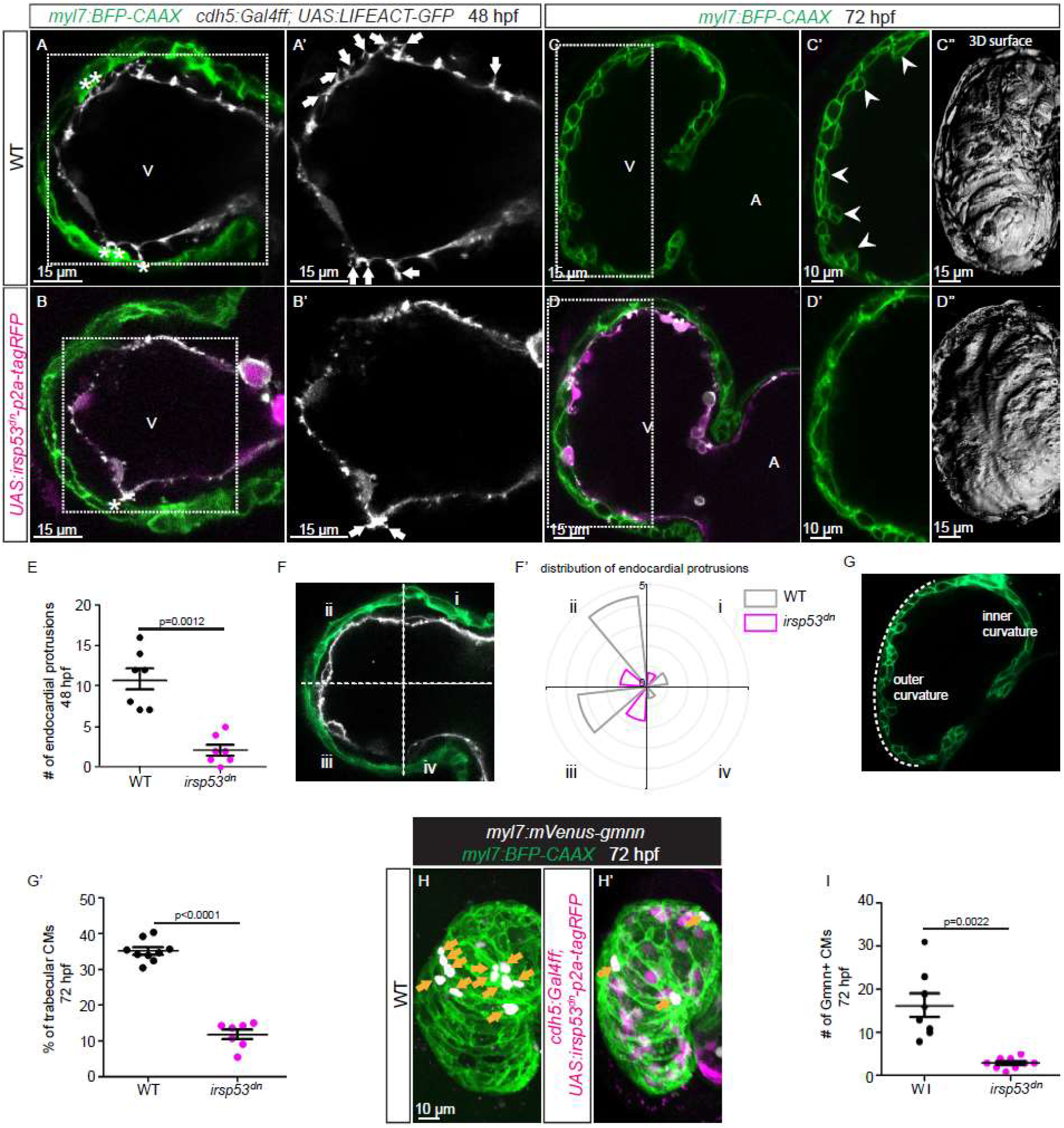
Blocking endocardial protrusion formation reduces cardiac trabeculation. **(A-D)** Confocal projection images of the heart of *Tg(myl7:BFP-CAAX)*; *Tg(cdh5:Gal4ff)*; *Tg(UAS:LIFEACT-GFP)*; +/- *Tg(UAS:irsp53^dn^-p2a-tagRFP)* zebrafish at 48 **(A-B)** and 72 **(C-D)** hpf. **(A-B)** Endocardial protrusions (white arrows) and touchdowns (white asterisks) are reduced in embryos with endothelial overexpression of *irsp53^dn^*. **(C-D)** Cardiac trabeculation (arrowheads) is reduced in larvae with endothelial overexpression of *irsp53^dn^*; **(C’’-D’’)** 3D rendering. **(E)** Quantification of the number of endocardial protrusions in wild-type and in embryos with endothelial overexpression of *irsp53^dn^* at 48 hpf. **(F-F’)** Illustration of the division of the 48 hpf ventricle into 4 regions **(F)**. Distribution and average number of endocardial protrusions in different regions of mid-sagittal sections of the ventricle from 48 hpf wild-type and *irsp53^dn^* embryos **(F’)**. **(G-G’)** Illustration of the division of the 72 hpf ventricle into the outer and inner curvature **(G)**. Quantification of the number of trabecular CMs in the outer curvature of wild-type and *irsp53^dn^* larvae at 72 hpf **(G’)**. **(H-H’)** 72 hpf larvae with endothelial overexpression of *irsp53^dn^* display a reduced number of *myl7*:mVenus-Gmnn^+^ CMs (yellow arrows) in their ventricle. **(I)** Quantification of the number of mVenus-Gmnn^+^ CMs in the ventricle of wild-type and *irsp53^dn^* larvae at 72 hpf. All images are ventral views, anterior to the top. V, ventricle; A, atrium. Data in graphs expressed as mean ± SEM.

### Apelin signaling positively regulates endocardial protrusion formation and myocardial trabeculation

We have recently shown that Apelin signaling regulates endothelial protrusion formation during angiogenesis in the zebrafish trunk (Helker et al., 2020). Therefore, we hypothesized that Apelin signaling might also regulate endocardial protrusion formation. To examine the expression pattern of the apelin ligand and receptor genes during heart development in zebrafish embryos, we first performed whole mount *in situ* hybridization. We detected *apln*, but no *apela*, expression within the heart (Figure 3-figure supplement 1A-D). For the receptor genes, we could only detect *aplnrb* expression in the heart (Figure 3-figure supplement 1E-H). In order to visualize the expressions of *apln* and *aplnrb* at single cell resolution in the heart, we examined the *TgBAC(apln:EGFP)* reporter line (Helker et al., 2020) line and generated a novel *Tg(aplnrb:VenusPEST)* reporter line. We detected apln:EGFP expression in the myocardium at 48 and 72 hpf (Figure 3A and B). Furthermore, we detected aplnrb:VenusPEST expression in the endocardium at 48 and 72 hpf (Figure 3C and D). These results suggest that *apln* is expressed in the myocardium while *aplnrb* is expressed in EdCs. Based on these results, we hypothesized that Apelin signaling plays a role during endocardium-myocardium interactions.

**Figure 3.**
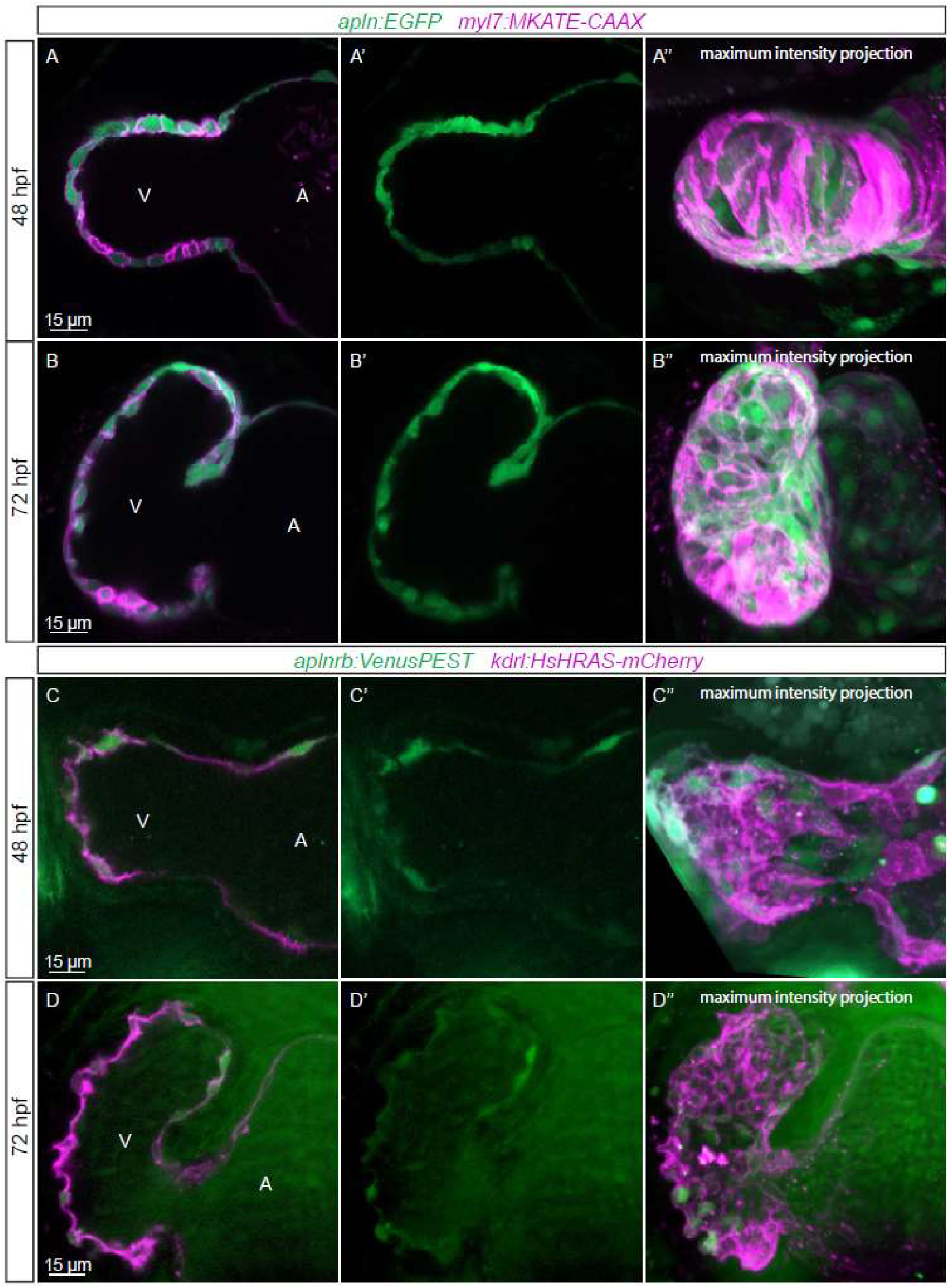
Expression pattern of Apelin signaling pathway components. **(A-D)** Confocal projection images of the heart of *TgBAC(apln:EGFP)*; *Tg(myl7:MKATE-CAAX)* **(A, B)** and *TgBAC(aplnrb:VenusPEST)*; *Tg(kdrl:HsHRAS-mCherry)* **(C, D)** zebrafish at 48 **(A, C)** and 72 **(B, D)** hpf. **(A’’-D’’)** Maximum intensity projections. **(A-B)** *TgBAC(apln:EGFP)* expression is detectable in the myocardium at 48 **(A)** and 72 **(B)** hpf. **(C-D)** *TgBAC(aplnrb:VenusPEST)* expression is detectable in the endocardium with higher expression in the ventricular endocardium at 48 **(C)** and 72 **(D)** hpf. All images are ventral views, anterior to the top. V, ventricle; A, atrium.

To test this hypothesis, we used mutants for *aplnra* (Helker et al., 2015), *aplnrb* (Helker et al., 2015), *apln* (Helker et al., 2015), and *apela* (Chng et al., 2013). Since *apela* mutants fail to form a heart (Chng et al., 2013), we did not analyze them. Most *aplnrb* mutants also fail to form a heart (Scott et al., 2007; Zeng et al., 2007), but some do (Figure 4-figure supplement 1C). By analyzing *aplnrb* mutants those do form a heart, we observed that they exhibit a reduced number of endocardial protrusions at 48 hpf (Figure 4-figure supplement 2A and B) and trabeculae at 72 hpf (Figure 4-figure supplement 2C and D). In wild-type embryos, the CJ between the endocardium and myocardium in the outer curvature of the ventricle appears to be mostly degraded at 72 hpf (Figure 4-figure supplement 2C); however, the CJ in *aplnrb* mutants appears to be thicker at this stage (Figure 4-figure supplement 2D). In addition, *aplnra* mutants exhibit a reduced number of trabeculae at 72 hpf (Figure 4-figure supplement 2E and F).

While *apln* mutants form a heart (Figure 4-figure supplement 1F), they display a significantly lower number of endocardial protrusions at 24 and 48 hpf (Figure 4A-D, G-I). In line with fewer endocardial protrusions, *apln* mutants exhibit a reduced number of endocardial touchdowns at 48 hpf (Figure 4C and D). Altogether, these results indicate that Apelin signaling regulates endocardial protrusion formation.

**Figure 4.**
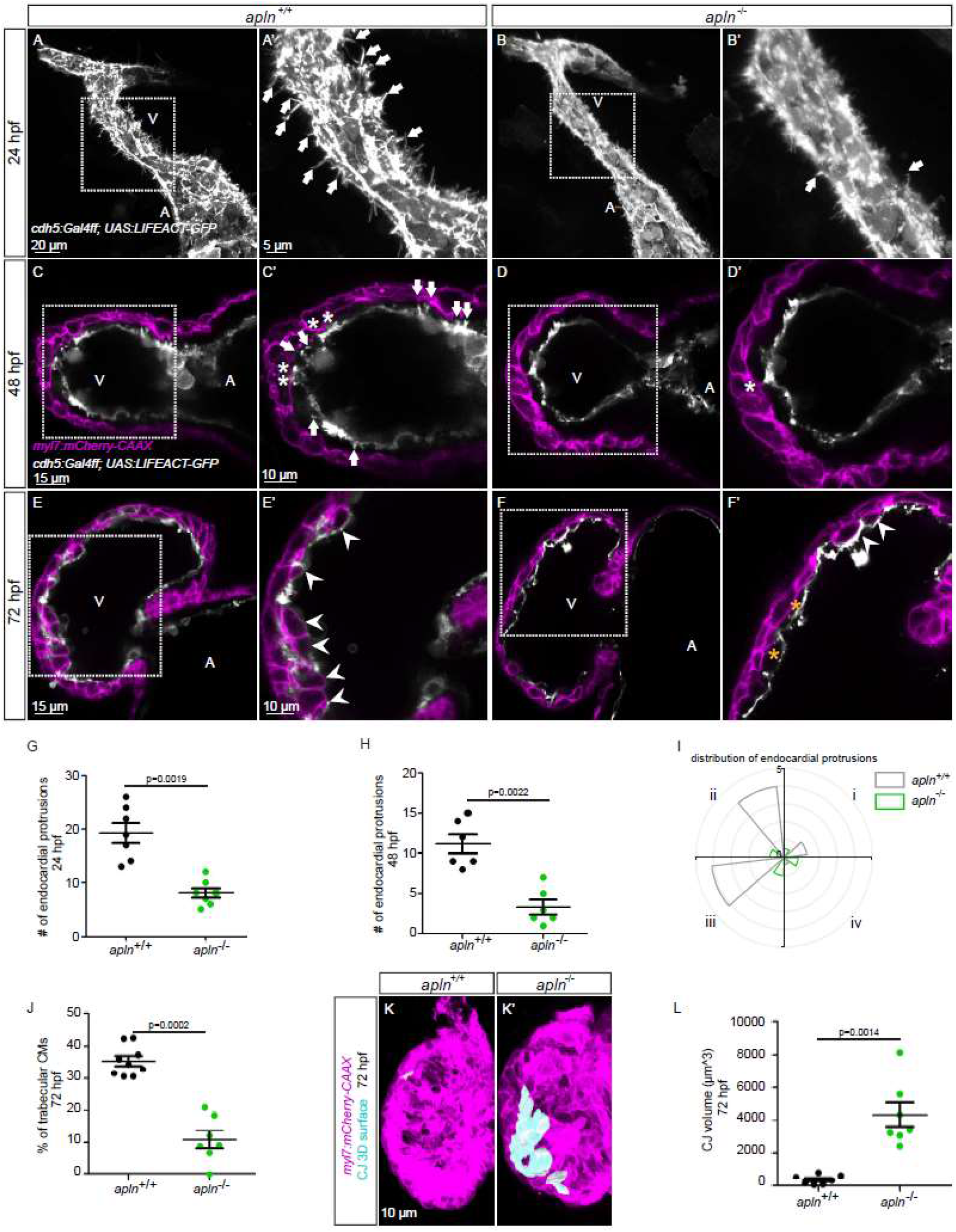
Loss of Apelin signaling leads to reduced endocardial protrusions and reduced myocardial trabeculation. **(A-F)** Confocal projection images of the heart of *Tg(cdh5:Gal4ff)*; *Tg(UAS:LIFEACT-GFP)* zebrafish at 24 hpf **(A-B)** and of the heart of *Tg(myl7:mCherry-CAAX)*; *Tg (cdh5:Gal4ff)*; *Tg(UAS:LIFEACT-GFP)* **(C-F)** zebrafish at 48 **(C-D)** and 72 **(E-F)** hpf. Maximum intensity projections **(A-B)** and mid-sagittal sections **(C-F)**. **(A)** Endocardial protrusions (arrows) in *apln^+/+^* embryos at 24 hpf. **(B)** Endocardial protrusions (arrows) are reduced in *apln^-/-^* siblings at 24 hpf. **(C-D)** Endocardial protrusions (arrows) and touchdowns (white asterisks) are reduced in *apln^-/-^* embryos **(D)** at 48 hpf compared with *apln^+/+^* siblings **(C)**. **(E-F)***apln^-/-^* larvae **(F)** exhibit reduced trabeculation (arrowheads) and thicker CJ (yellow asterisks) at 72 hpf compared with *apln^+/+^* siblings **(E)**. **(G-H)** Quantification of the number of endocardial protrusions in the ventricle of *apln^+/+^* and *apln^-/-^* siblings at 24 **(G)** and 48 **(H)** hpf. **(I)** Distribution and average number of endocardial protrusions in different regions of mid-sagittal sections of the ventricle from 48 hpf *apln^+/+^* and *apln^-/-^* siblings. **(J)** Quantification of the number of trabecular CMs in the outer curvature of *apln^+/+^* and *apln^-/-^* siblings at 72 hpf. **(K-K’)** Maximum intensity projections. *apln^-/-^* larvae **(K’)** exhibit a thicker CJ at 72 hpf compared with *apln^+/+^* siblings **(K). (L)** Quantification of the CJ volume in the outer curvature of *apln^+/+^* and *apln^-/-^* siblings at 72 hpf. All images are ventral views, anterior to the top. V, ventricle; A, atrium; *+/+*, *apln^+/+^*; *-/-, apln^-/-^*. Data in graphs expressed as mean ± SEM.

To examine the function of Apelin dependent endocardial protrusions on cardiac trabeculation, we first analyzed trabecular formation in *apln* mutants. Homozygous *apln* mutants exhibit a reduced number of trabeculae at 72 hpf (Figure 4E, F, and J). In order to analyze CM proliferation, we performed EdU labeling and quantified EdU^+^ CMs in *apln* mutant and wild-type sibling larvae. Homozygous *apln* mutants exhibit a significantly decreased number of EdU^+^ CMs in their ventricle (Figure 4-figure supplement 3). In addition, *apln* mutants display a significantly thicker CJ compared with wild-type siblings at 72 hpf (Figure 4C-F, K, and L). However, we did not observe any obvious defects in sarcomere formation (Figure 4-figure supplement 4) in *apln* mutants at 72 hpf.

Notch signaling negatively regulates endothelial sprouting and protrusion formation in several vascular beds (Hellstrom et al., 2007; Leslie et al., 2007; Siekmann and Lawson, 2007; Suchting et al., 2007). In order to analyze whether Notch signaling also regulates endocardial protrusion formation, we blocked Notch signaling by treating embryos with the γ-secretase inhibitor RO4929097, and observed a decrease of Notch reporter expression (Figure 4-figure supplement 5A and B) as well as an increased number of endocardial protrusions in the ventricle (Figure 4-figure supplement 5C-E). Together, these results show that myocardial derived Apelin positively regulates endocardial protrusion formation while Notch signaling negatively regulates it. Furthermore, Apelin signaling is also required for cardiac trabeculation, possibly via the formation of endocardial protrusions.

### The effect of endocardial *nrg2a* in trabeculation is mediated by endocardial protrusions

Nrg-ErbB signaling is indispensable for cardiac trabeculation in mouse and zebrafish (Gassmann et al., 1995; Lee et al., 1995; Meyer and Birchmeier, 1995; Lai et al., 2010; Liu et al., 2010; Rasouli and Stainier, 2017). To determine whether endocardial protrusions modulate Nrg-ErbB signaling, we overexpressed *nrg2a* in the endocardium using a *Tg(fli1a:nrg2a-p2a-tdTomato)* line (Rasouli and Stainier, 2017). Overexpression of *nrg2a* in the endocardium results in hypertrabeculation as well as a multilayered myocardium (Figure 5A, B, and E-G). Strikingly, overexpressing *nrg2a* in the endothelium while blocking endocardial protrusion formation by endothelial overexpression of *irsp53^dn^* is not sufficient to restore cardiac trabeculation and induce CM multilayering (Figure 5C-C’’’ and E-G). In line with these results, overexpressing *nrg2a* in the endothelium of homozygous *apln* mutants is not sufficient to restore cardiac trabeculation and induce CM multilayering (Figure 5D-D’’’ and E-G). Importantly, we did not detect a change in the expression levels of *nrg2a* in *apln* mutant hearts at 48 hpf (Figure 5-figure supplement 1). Taken together, these data suggest that endocardial protrusions are required for Nrg-ErbB signaling.

**Figure 5.**
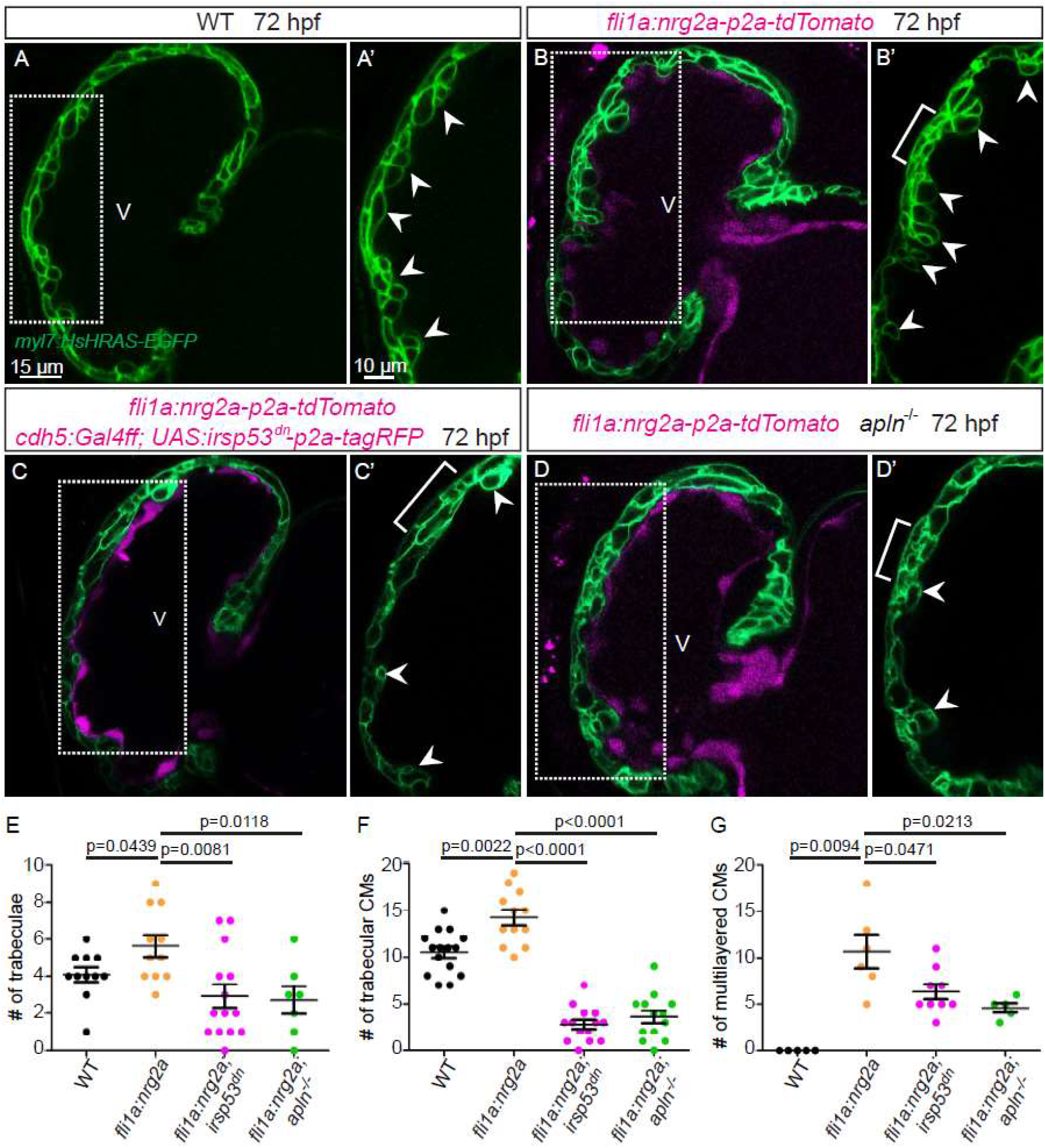
Endocardial protrusions are necessary for *nrg2a* overexpression phenotypes. **(A-D)** Confocal projection images of the heart of *Tg(myl7:HsHRAS-EGFP)* larvae at 72 hpf. **(A-B)** Overexpression of *nrg2a* in the endothelium **(B)** leads to an increased number of trabeculae (arrowheads) and the multilayering of CMs (brackets) compared with wild-type **(A)**. **(C)** Larvae with endothelial overexpression of *nrg2a* and *irsp53^dn^* exhibit a reduced number of trabeculae (arrowheads) and of multilayered CMs (brackets) compared with larvae with endothelial overexpression of *nrg2a* alone **(B)**. **(D)** *apln* mutant larvae with endothelial overexpression of *nrg2a* exhibit a reduced number of trabeculae (arrowheads) and of multilayered CMs (brackets) compared with larvae with endothelial overexpression of *nrg2a* alone **(B)**. **(E)** Quantification of the number of trabeculae. **(F)** Quantification of the number of trabecular CMs. **(G)** Quantification of the number of multilayered CMs in the ventricle. Brackets indicate multilayered CMs. All images are ventral views, anterior to the top. V, ventricle. Data in graphs expressed as mean ± SEM.

### Genetically blocking endocardial protrusion formation attenuates Erk signaling in cardiomyocytes

An important molecule in the Nrg/ErbB signaling pathway is the extracellular signal-regulated kinase Erk (Lai et al., 2010). In order to visualize Erk activity in CMs in living zebrafish, as a readout of ErbB signaling, we generated novel reporter lines (*Tg(myl7:ERK-KTR-Clover-p2a-H2B-tagBFP)* and *Tg(myl7:ERK-KTR-Clover-p2a-H2B-mScarlet)*) by using the kinase translocation reporter (KTR) technology (Regot et al., 2014; de la Cova et al., 2017). When Erk is inactive, the KTR is unphosphorylated and Clover can be detected in the nucleus; in contrast, when Erk is active, the KTR is phosphorylated and Clover can be detected in the cytoplasm (de la Cova et al., 2017). We observed that most ventricular CMs in wild-type larvae display active Erk signaling with cytoplasmic Clover expression (Figure 6A). Treating the reporter with a MEK inhibitor led to an increased number of ventricular CMs with nuclear Clover expression (i.e., inactive Erk signaling) indicating that our reporter is functional (Figure 6-figure supplement 1). Next, we treated this reporter with an ErbB2 inhibitor and found an increased number of ventricular CMs with nuclear Clover expression (Figure 6B). To determine whether endocardial protrusions modulate myocardial Erk signaling activity, we genetically blocked endocardial protrusions via endothelial overexpression of *irsp53^dn^* (Figure 6C). We observed more ventricular CMs with nuclear Clover expression in the larvae overexpressing *irsp53^dn^* (Figure 6C and E) compared with control (Figure 6A and E), indicating more ventricular CMs with inactive Erk signaling. In line with these results, we observed more CMs with inactive Erk signaling in homozygous *apln* mutants (Figure 6D and E) compared with wild-type siblings (Figure 6A and E). Altogether, these observations indicate that Apelin signaling dependent endocardial protrusions modulate Nrg/ErbB/Erk signaling in CMs.

**Figure 6.**
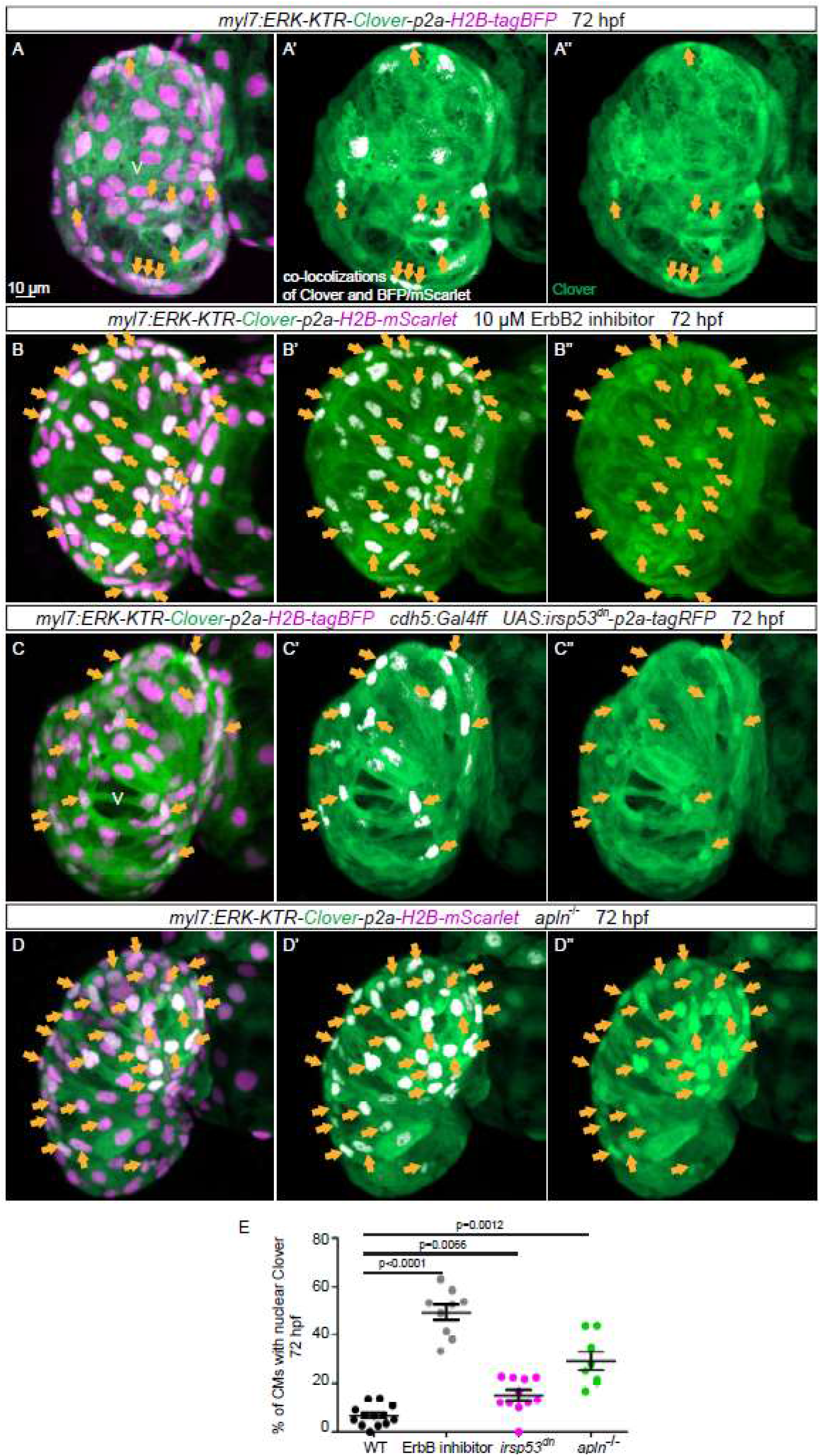
Blocking endocardial protrusion formation reduces myocardial Erk signaling activity. **(A-D)** Maximum intensity projections of confocal images of the heart of *Tg(myl7:ERK-KTR-Clover-p2a-H2B-tagBFP/mScarlet)* larvae at 72 hpf. **(A)** Visualization of Erk activity by a CM specific ERK-KTR reporter. Nuclear Clover expression (arrows) indicates CMs with inactive Erk signaling. **(B)** Larvae treated with an ErbB2 inhibitor exhibit an increased number of CMs with inactive Erk signaling (arrows) compared with control larvae **(A)**. **(C)** Larvae with endothelial overexpression of *irsp53^dn^* exhibit an increased number of CMs with inactive Erk signaling (arrows) compared with wild-type larvae **(A)**. **(D)** *apln* mutant larvae exhibit an increased number of CMs with inactive Erk signaling (arrows) compared with *apln^+/+^* siblings. **(E)** Quantification of ventricular CMs with nuclear Clover expression. All images are ventral views, anterior to the top. V, ventricle. Data in graphs expressed as mean ± SEM.

## Discussion

### Endocardial protrusions contribute to trabeculation

Cardiac trabeculation is initiated, at least in zebrafish, by individual CMs delaminating from the compact myocardial wall and protruding into the lumen (Liu et al., 2010; Staudt et al., 2014; Jimenez-Amilburu et al., 2016; Priya et al., 2020). Several studies have reported that the endocardium plays an important role during cardiac trabeculation (Grego-Bessa et al., 2007; Lai et al., 2010; D’Amato et al., 2016; Rasouli and Stainier, 2017; Del Monte-Nieto et al., 2018; Qu et al., 2019). Furthermore, it has recently been shown that EdCs, similar to ECs, undergo sprouting (Del Monte-Nieto et al., 2018). However, in comparison with endocardial sprouting, little is known about the morphogenetic events underlying endocardial sprouting and their effect on cardiac trabeculation.

In mouse, endocardial sprouting and touchdown formation occur early during cardiac trabeculation (Del Monte-Nieto et al., 2018). These observations are in line with our data in zebrafish suggesting that the morphogenetic events of cardiac trabeculation are evolutionarily conserved. CM delamination and trabeculation occur in the outer curvature of the ventricle (Liu et al., 2010; Jimenez-Amilburu et al., 2016; Rasouli and Stainier, 2017). This observation is in line with our finding that endocardial protrusions are mostly located in the outer curvature of the ventricle. The spatial and temporal correlation between the emergence of endocardial protrusions and CM delamination therefore suggests a role for endocardial protrusions in cardiac trabeculation.

### Molecular regulators of endocardial sprouting

During sprouting angiogenesis, so-called tip cells lead the new sprouts (Gerhardt, 2008). Tip cells dynamically extend filopodia to identify growth factors in their environment (Gerhardt, 2008). Apelin and Notch signaling have been previously identified as regulators of endothelial filopodia formation (Hellstrom et al., 2007; Suchting et al., 2007; Helker et al., 2020). In contrast, the pathways regulating endocardial sprouting are largely unknown. Only Tie2 signaling has been identified to date as a regulator of endocardial sprouting, and Tie2 deficient mice exhibit fewer endocardial touchdowns (Qu et al., 2019). We have recently shown that Apelin signaling regulates filopodia formation during sprouting angiogenesis in the trunk (Helker et al., 2020). In line with these published observations, we now show that Apelin regulates endocardial filopodia formation and endocardial sprouting (Figure 6-figure supplement 2), highlighting a conserved role for Apelin signaling during endothelial and endocardial sprouting.

Consistent with the regulation of sprouting angiogenesis by Notch signaling in ECs (Hellstrom et al., 2007; Leslie et al., 2007; Siekmann and Lawson, 2007; Suchting et al., 2007), we found that Notch signaling also negatively regulates endocardial protrusion formation. Interestingly, inhibition of Notch signaling also leads to an increased number of delaminated CMs and trabeculae (Han et al., 2016; Priya et al., 2020).

### Endocardium-myocardium communication is essential for trabeculation

Paracrine communication is usually thought to be based on the diffusion of soluble morphogens. The Nrg/ErbB signaling pathway, which is required for cardiac trabeculation, resembles such a classical paracrine signaling pathway (Gassmann et al., 1995; Lee et al., 1995; Meyer and Birchmeier, 1995; Lai et al., 2010; Liu et al., 2010; Rasouli and Stainier, 2017). Several studies have shown that endocardial derived Nrg is required to activate ErbB receptor complexes on CMs (Gassmann et al., 1995; Meyer and Birchmeier, 1995; Grego-Bessa et al., 2007; Rasouli and Stainier, 2017).

Like other receptor tyrosine kinases, ErbB receptors activate multiple signaling cascades, including the MAPK cascade, upon ligand stimulation, leading to the phosphorylation of ERK1/2 (Sweeney et al., 2001; Wee and Wang, 2017). Accordingly, attenuated phosphorylation of ERK in CMs is observed in mice deficient in Nrg1/ErbB signaling (Lai et al., 2010). By analyzing a novel reporter of Erk activity in CMs, we observed that the inhibition of endocardial protrusions as well as the genetic inactivation of Apelin signaling lead to attenuated Erk phosphorylation in CMs. Together, these data suggest that Apelin signaling dependent endocardial protrusions modulate ErbB signaling in CMs (Figure 6-figure supplement 2).

It has recently been shown that filopodia from ECs modulate neurogenesis by affecting progenitor cell proliferation in the developing brain of mice and zebrafish (Di Marco et al., 2020; Taberner et al., 2020). Of interest, ErbB signaling is also known for its function within the nervous system (Buonanno and Fischbach, 2001). Thus, one might speculate that Nrg/ErbB signaling also plays a role during the modulation of neurogenesis by endothelial filopodia. Several studies reported cell to cell communication by cytonemes in different animal models (Ramirez-Weber and Kornberg, 1999; Holzer et al., 2012; Luz et al., 2014; Sagar et al., 2015). Whether endocardial protrusions qualify as cytonemes needs further analysis. However, our data indicate that Apelin dependent endocardial protrusions are required for the communication between endocardial and myocardial cells via Nrg/ErbB signaling (Figure 6-figure supplement 2).

In summary, our work describes how endocardial sprouting is integrated into Nrg/ErbB signaling and cardiac trabeculation. Furthermore, we identify Apelin signaling as a regulator of endocardial sprouting.

## Acknowledgements

We thank Gisela Thana Hartmann, Sarah Howard, Dr. Radhan Ramadass, and all fish facility staff for their technical support; Dr. Thomas Juan, Dr. Samuel Capon, Dr. Jordan Welker, Giulia Boezio and Yiu Chun Law for critical comments on the manuscript; Dr. Stefan Baumeister for the schematic model; and Dr. Gonzalo del Monte-Nieto for discussion. Research in the Stainier laboratory is supported in part by the Max Planck Society, the DFG (SFB 834/4) and the Leducq Foundation. Research in the Helker laboratory is supported in part by the DFG (SFB 834/4).

## Author contributions

J.Q., D.Y.R.S. and C.S.M.H. designed experiments, J.Q. and A.R. performed experiments, A.R., R.P., S.M. provided unpublished transgenic lines, J.Q., D.Y.R.S., C.S.M.H. analyzed data, J.Q., D.Y.R.S., and C.S.M.H. wrote the manuscript. All authors commented on the manuscript.

## Author information

The authors declare no competing interests.

## Supplemental videos

**Figure 1-video 1. Endocardial touchdowns during cardiac contraction. Related to figure 1E–1H.**

Beating 48 hpf zebrafish heart. Magenta, myocardium; white, endocardium.

**Figure 1-video 2. Endocardial protrusions extend along delaminating CMs at 60 hpf. Related to figure 1C’’’.**

3D surface rendering of a 60 hpf ventricle. Magenta, myocardium; white, endocardium; yellow, endocardial protrusions extending along delaminating CMs.

**Figure 1-video 3. Endocardial protrusions are in close proximity to trabecular CMs at 72 hpf. Related to figure 1D’’’.**

3D surface rendering of a 72 hpf ventricle. Magenta, myocardium; white, endocardium; yellow, endocardial protrusions in close proximity to trabecular CMs.

---

**Figure 1-figure supplement 1.**
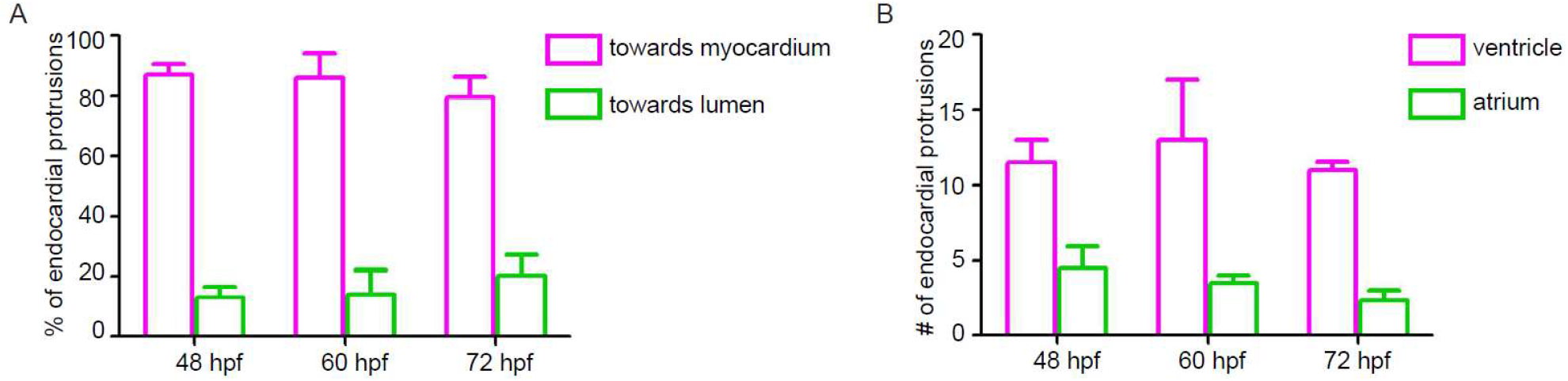
Endocardial protrusions in the ventricle extend mainly towards the myocardium. **(A)** Quantification of the direction of endocardial protrusions; most endocardial protrusions extend towards the myocardium. **(B)** Quantification of the average number of endocardial protrusions in the ventricle and atrium. n=9 in each group.

**Figure 1-figure supplement 2.**
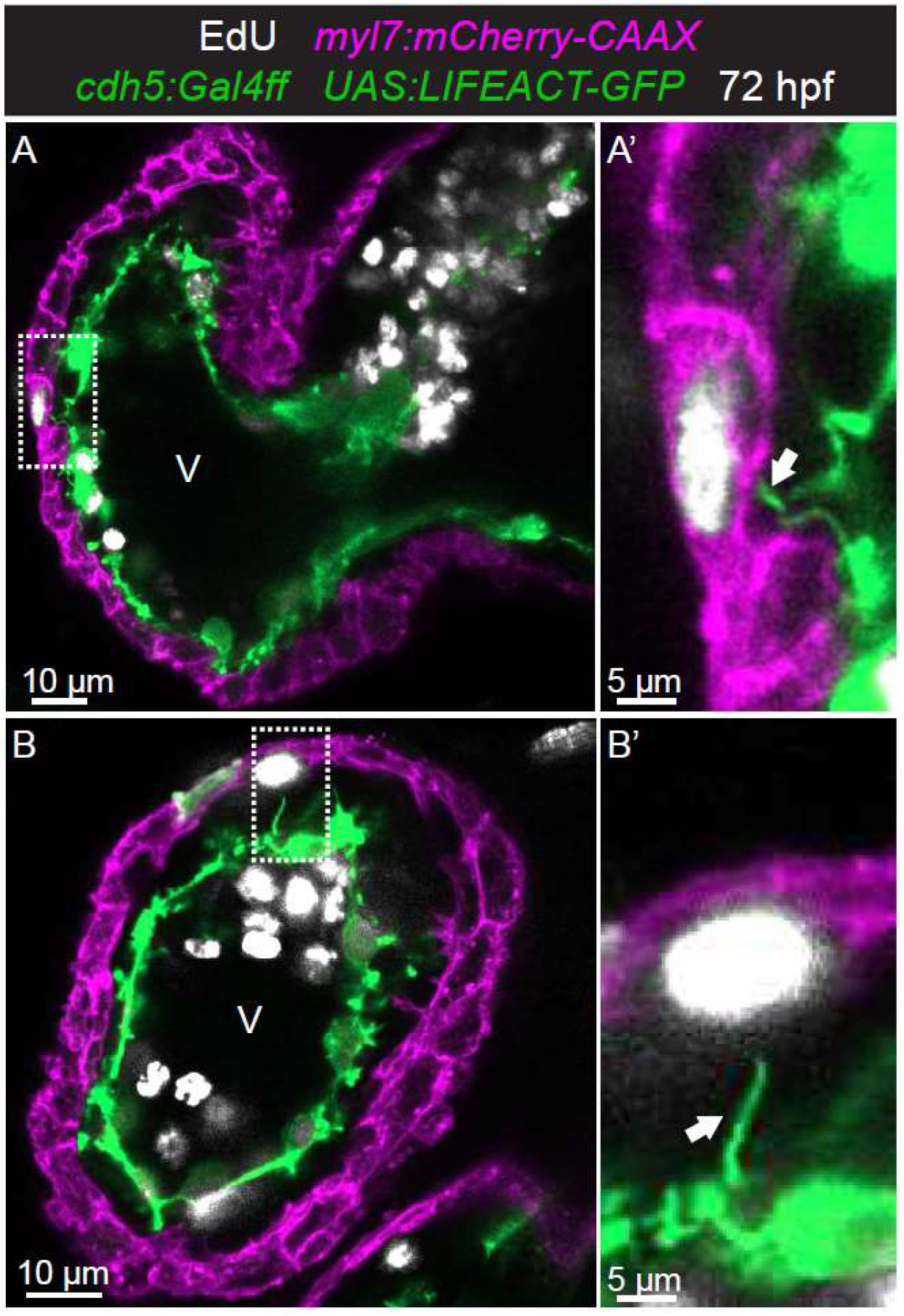
Endocardial protrusions most often contact proliferating CMs. **(A-B)** Confocal projection images of 72 hpf *Tg(myl7:mCherry-CAAX)*; *Tg (cdh5:Gal4ff)*; *Tg(UAS:LIFEACT-GFP)* hearts after EdU labeling from 28 to 72 hpf. **(A’-B’)** Arrows point to endocardial protrusions close to EdU^+^ CMs (n=8). All images are ventral views, anterior to the top. V, ventricle.

**Figure 3-figure supplement 1.**
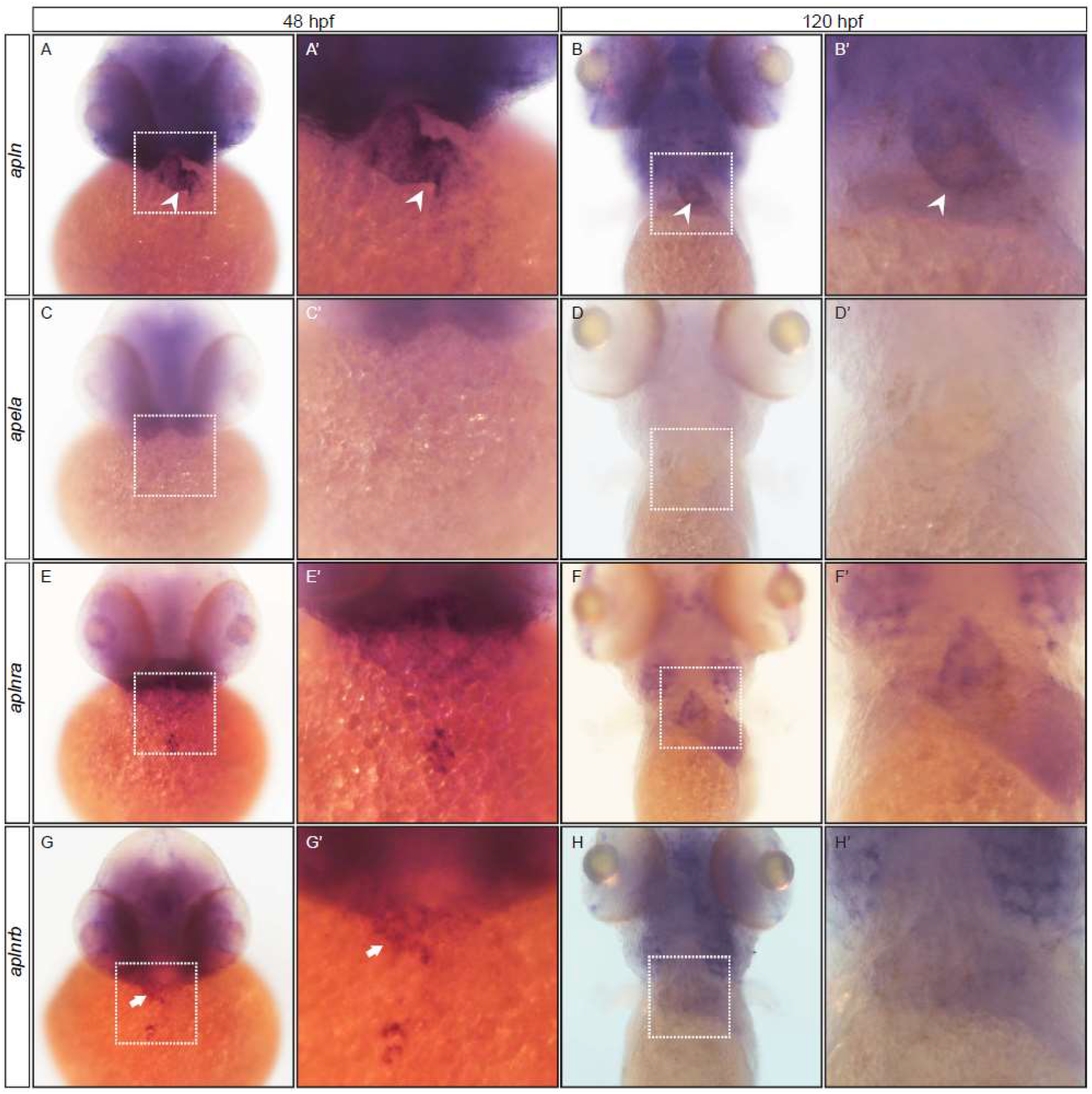
Expression of Apelin signaling ligand and receptor genes by *in situ* hybridization. **(A-H)** Expression of Apelin signaling ligand and receptor genes at 48 and 120 hpf. **(A-B)** *apln* is expressed in the developing heart (arrowheads). **(C-D)** *apela* expression is not detected in the developing heart. **(E-F)** *aplnra* expression is not detected in the developing heart. **(G-H)** *aplnrb* is expressed in the developing heart (arrows). White box, heart region.

**Figure 4-figure supplement 1.**
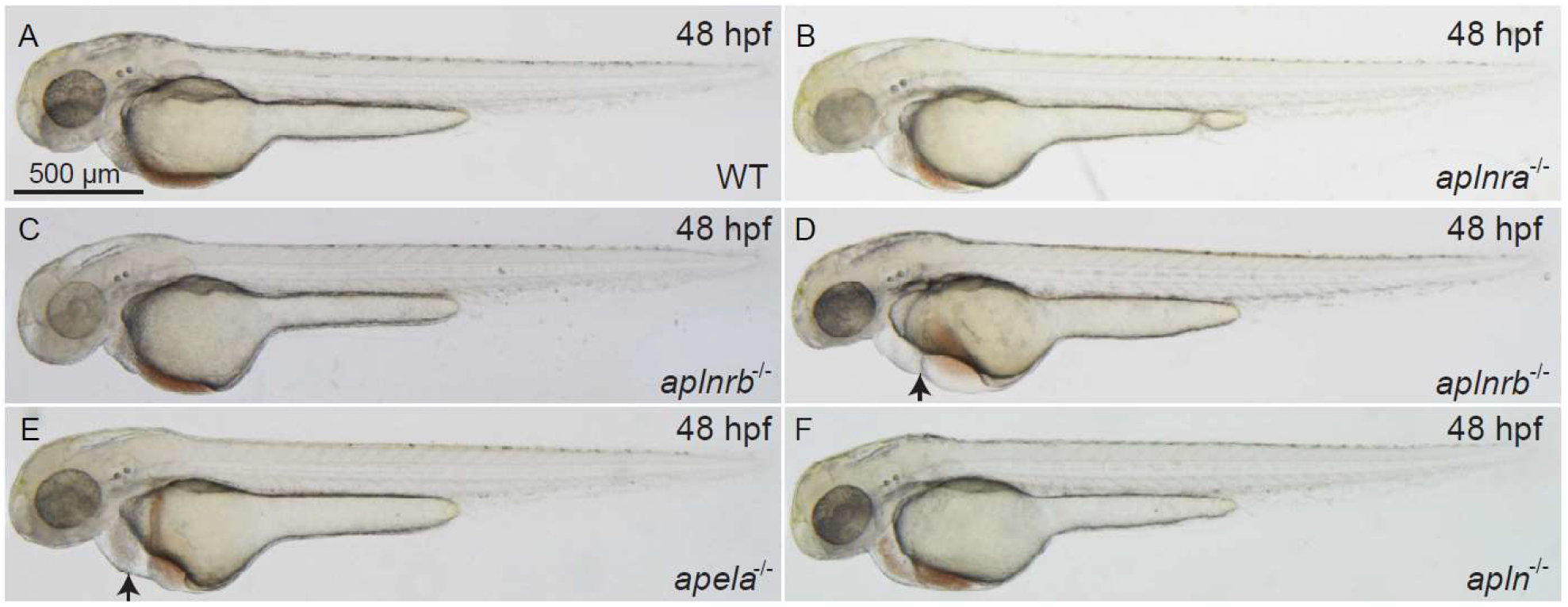
Bright field pictures of *aplnra, aplnrb, apela* and *apln* mutants. **(A-F)** Brightfield pictures (lateral views) of 48 hpf wild-type **(A)**, *aplnra* mutant **(B)**, *aplnrb* mutant without pericardial edema **(C)**, *aplnrb* mutant with pericardial edema (arrow) **(D)**, *apela* mutant with pericardial edema (arrow) **(E)**, and *apln* mutant without pericardial edema **(F)**.

**Figure 4-figure supplement 2.**
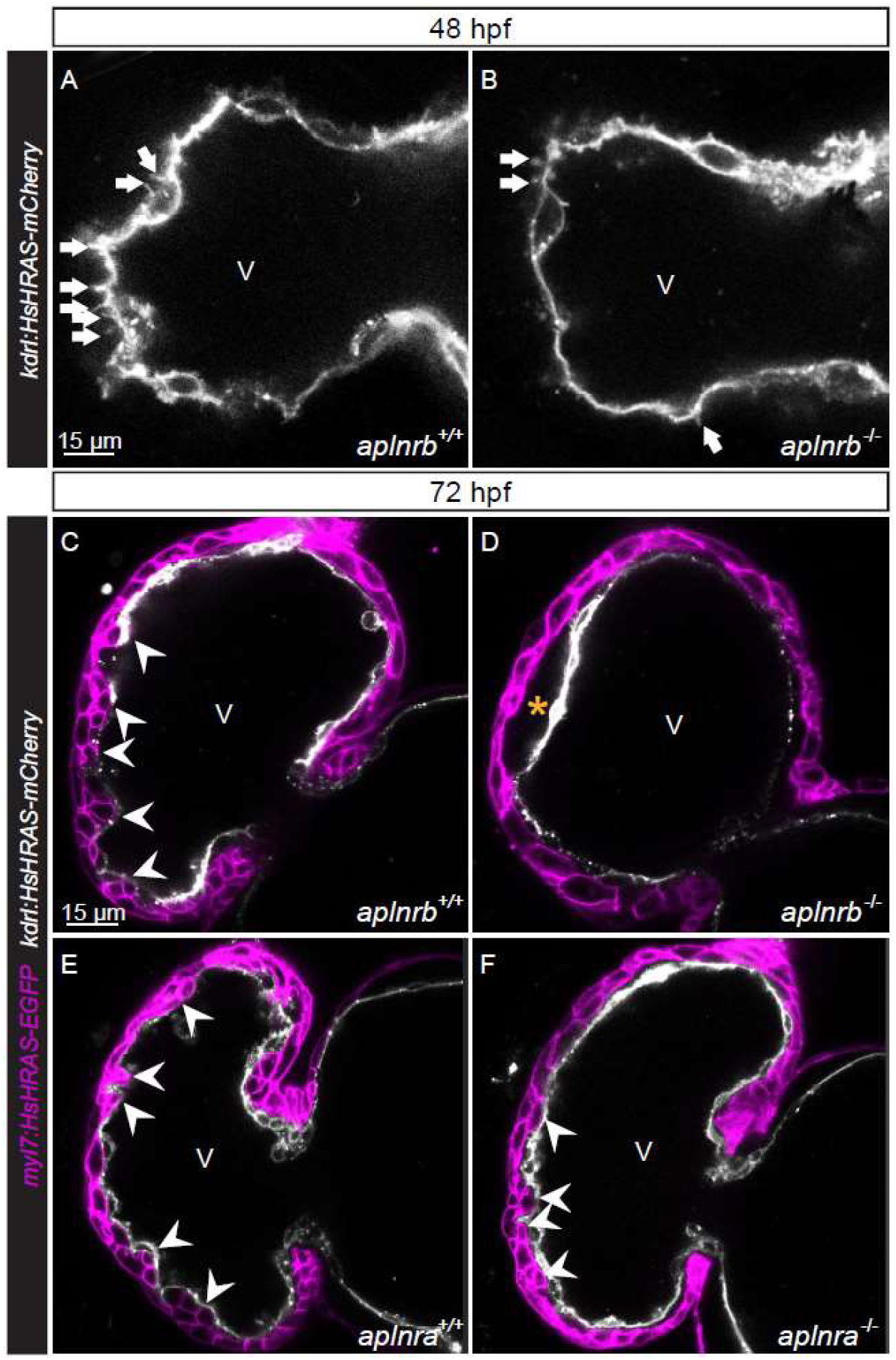
*aplnrb* mutants exhibit reduced endocardial protrusion formation and trabeculation and *aplnra* mutant exhibit a mild reduction in trabeculation. **(A–F)** Confocal projection images of the heart of *Tg(kdrl:HsHRAS-mCherry)* **(A-B)** and *Tg(myl7:HsHRAS-EGFP)*; *Tg(kdrl:HsHRAS-mCherry)* **(C-F)** zebrafish at 48 **(A-B)** and 72 **(C-F)** hpf. **(A–B)***aplnrb^-/-^* embryos exhibit fewer endocardial protrusions (arrows) compared with *aplnrb^+/+^* siblings **(A)** at 48 hpf. **(C-D)***aplnrb^-/-^* larvae **(D)** exhibit reduced trabeculation (arrowheads) and thicker CJ (asterisk) compared with *aplnrb^+/+^* siblings **(C)** at 72 hpf. **(E-F)***aplnra^-/-^* larvae **(F)** exhibit a mild reduction of trabeculation compared with *aplnra^+/+^*siblings **(E)** at 72 hpf. V, ventricle.

**Figure 4-figure supplement 3.**
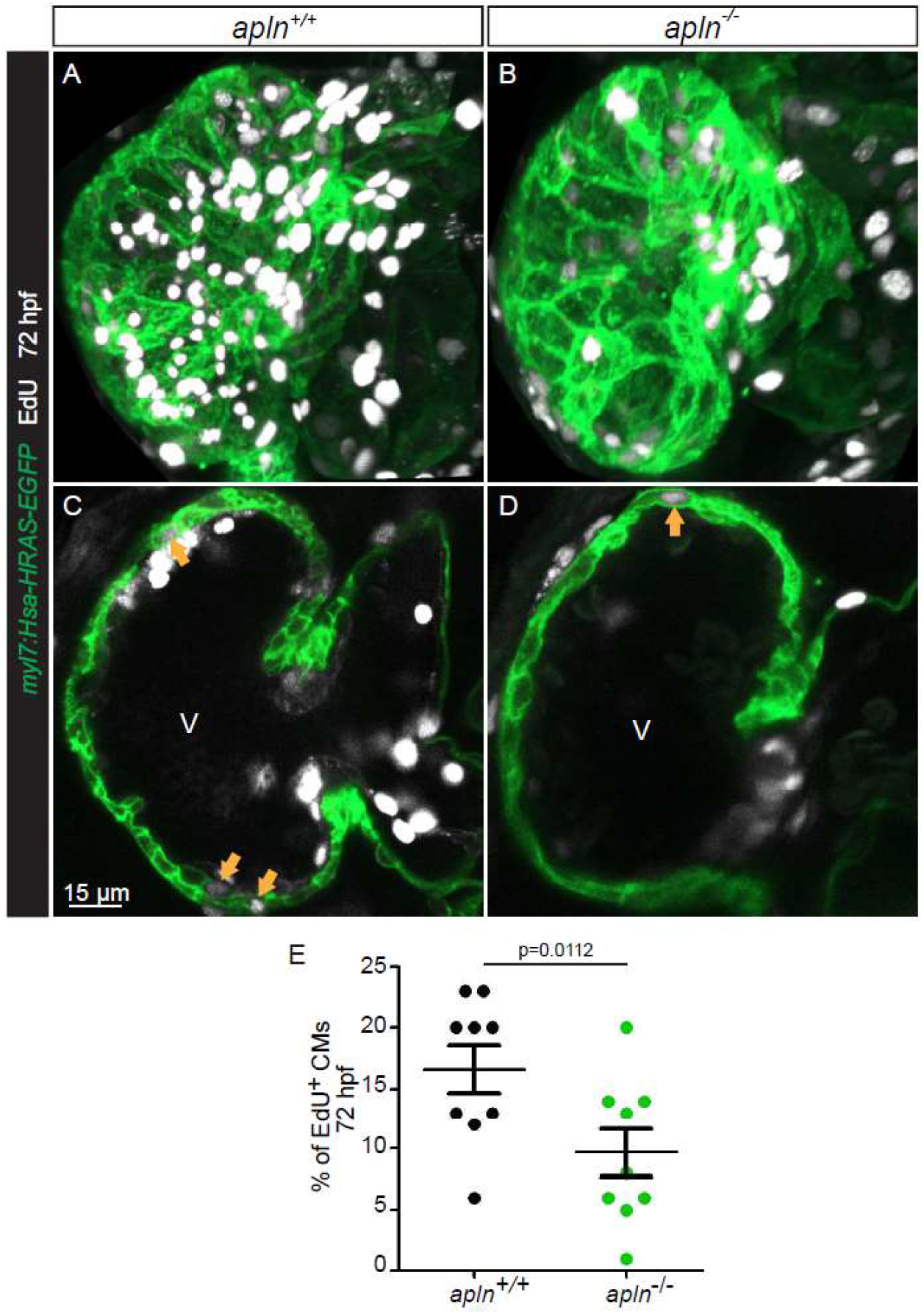
Apelin signaling regulates CM proliferation in the ventricle. **(A-D)** Confocal projection images of the heart of *Tg(myl7:HsHRAS-EGFP)* larvae at 72 hpf. **(A-B)** Maximum intensity projections of confocal images. **(C-D)** Mid-sagittal sections of A and B, respectively. *apln^-/-^* larvae **(D)** exhibit fewer proliferating CMs (arrows) in the ventricle compared with *apln^+/+^* siblings **(C)**. **(E)** Quantification of EdU^+^ CMs in the ventricle of *apln^+/+^* and *apln^-/-^* siblings. V, ventricle. Data in graphs expressed as mean ± SEM.

**Figure 4-figure supplement 4.**
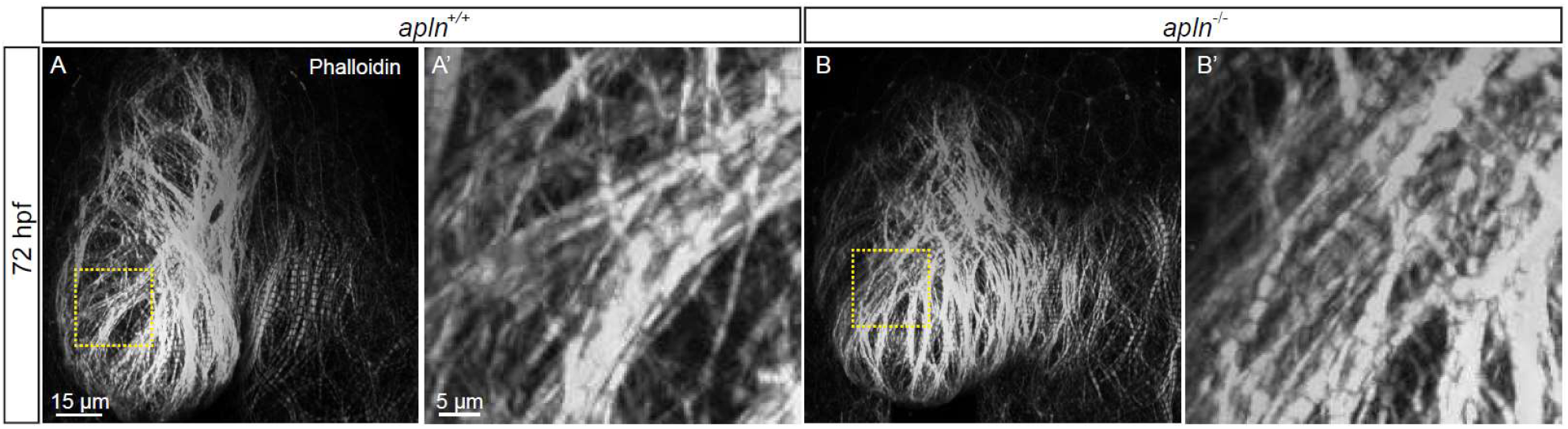
Wild-type like heart function and sarcomere structure in *apln^-/-^* larvae. **(A-B)** Quantification of heart rate **(A)** and ejection fraction **(B)** of *apln^+/+^* and *apln^-/-^* siblings. **(C-D)** Confocal projection images. Maximum intensity projections of confocal images of the heart of 72 hpf larvae stained with Phalloidin. Sarcomere formation does not appear to be affected in *apln^-/-^* larvae **(C)** compared with *apln^+/+^*siblings **(D)** (*apln^+/+^*, n=5; *apln^-/-^*, n=4). V, ventricle.

**Figure 4-figure supplement 5.**
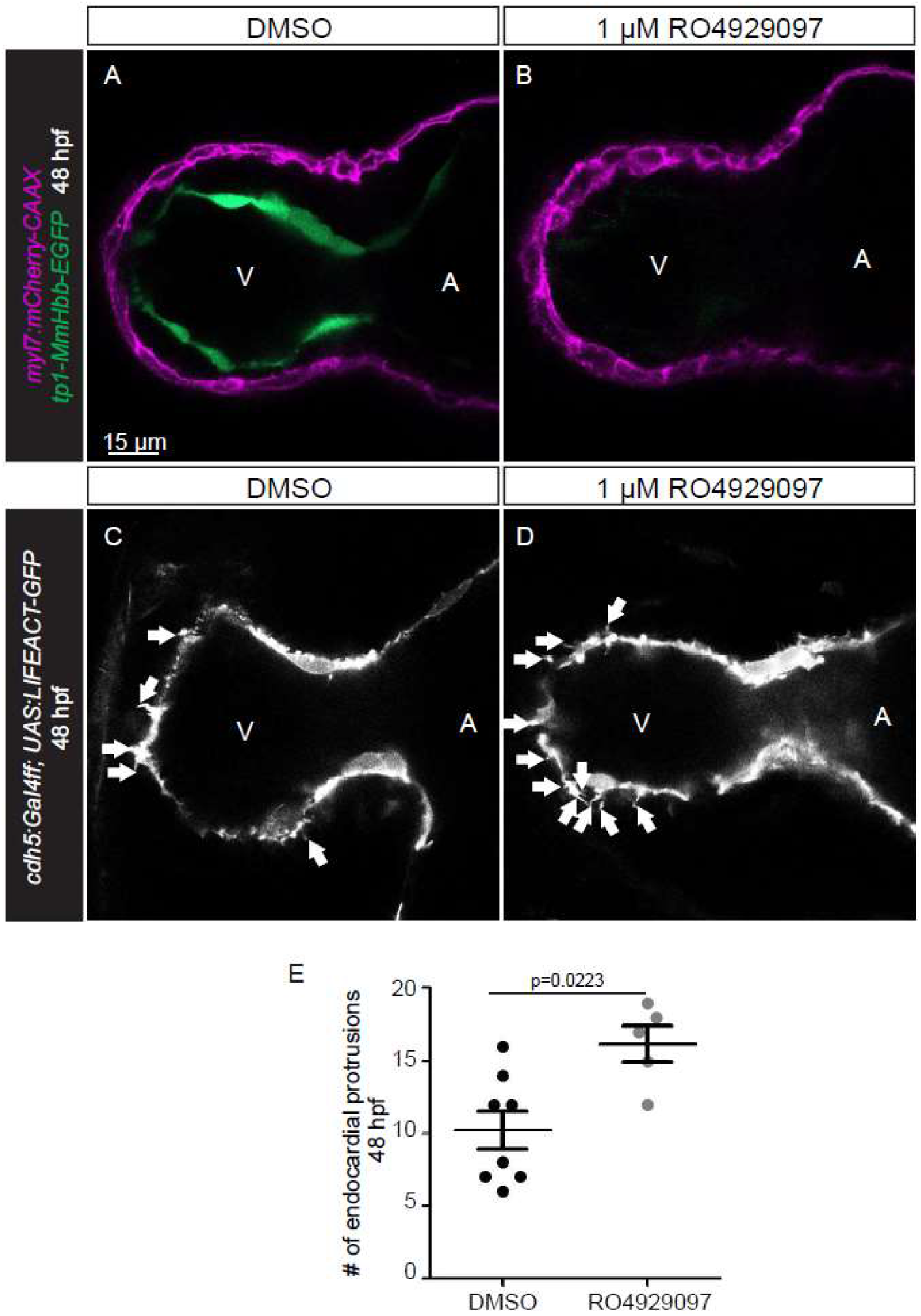
Notch signaling represses endocardial protrusion formation. **(A-D)** Confocal projection images of the heart of *Tg(myl7:mCherry-CAAX)*; *Tg(tp1-MmHbb:EGFP)* **(A-B)** and *Tg(cdh5:Gal4ff)*; *Tg(UAS:LIFEACT-GFP)* **(C-D)** embryos at 48 hpf. **(A-B)** Treatment with 1 μM of the Notch inhibitor RO4929097 from 24 to 48 hpf blocks the expression of the *Tg(tp1-MmHbb:EGFP)* Notch reporter in the endocardium. **(C-D)** Embryos treated with the Notch inhibitor exhibit more endocardial protrusions (arrows). **(E)** Quantification of the number of endocardial protrusions in the ventricle of DMSO and RO4929097 treated embryos at 48 hpf. All images are ventral views, anterior to the top. V, ventricle; A, atrium. Data in graphs expressed as mean ± SEM.

**Figure 5-figure supplement 1.**
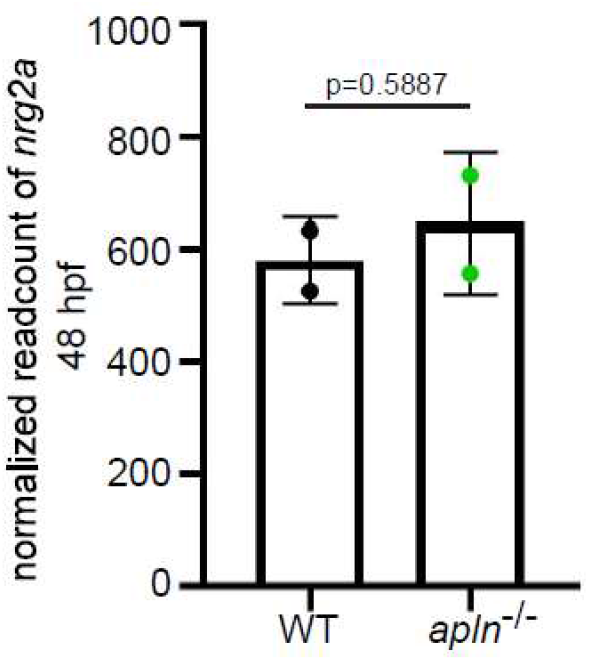
*nrg2a* expression does not appear to be affected in *apln* mutants. *nrg2a* mRNA levels in extracted hearts from wild types and *apln* mutants at 48 hpf (from RNA-seq). Data in graphs expressed as mean ± SEM.

**Figure 6-figure supplement 1.**
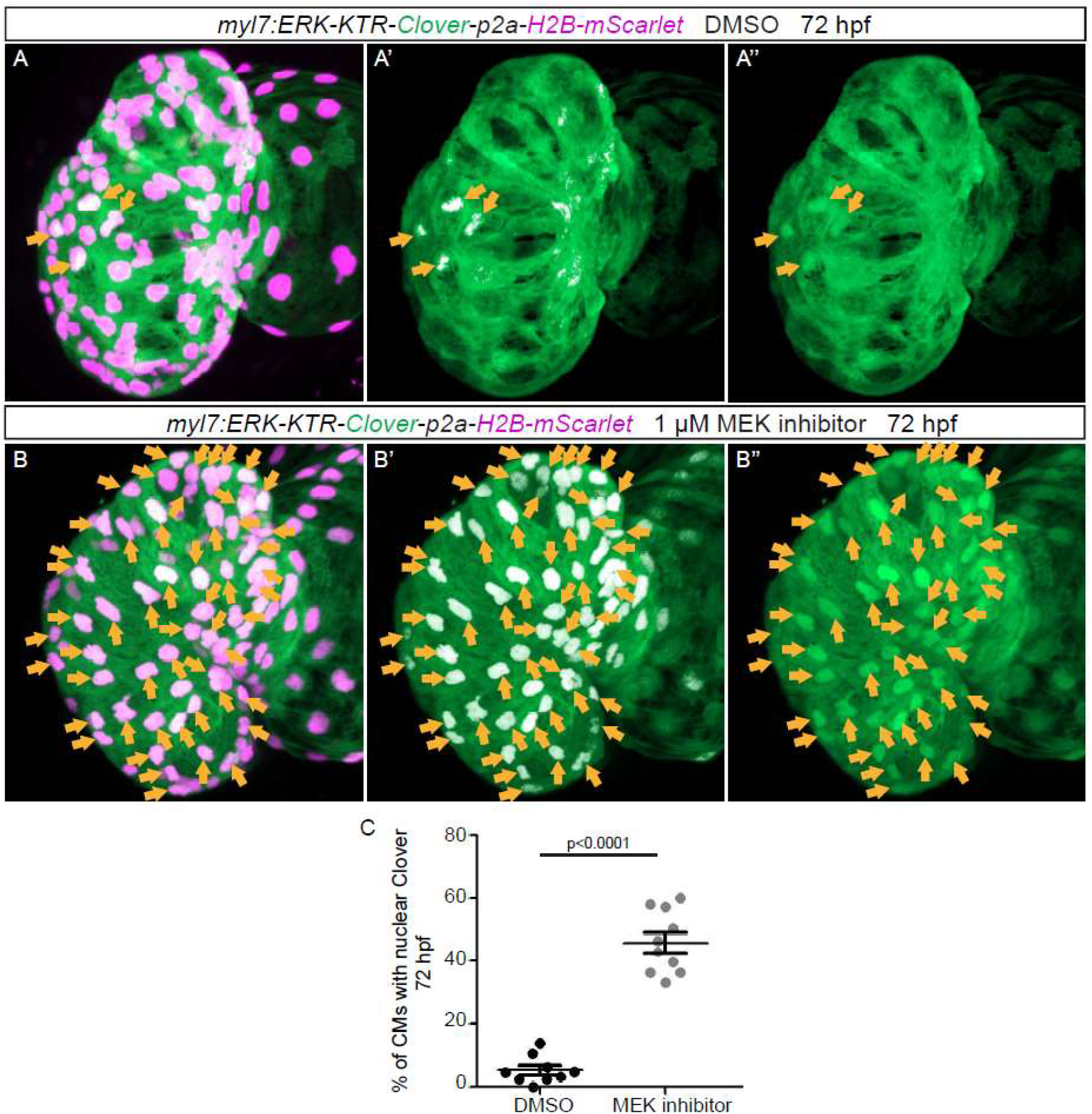
Erk inhibitor represses the activity of Erk in the Erk reporter line. **(A-B)** Confocal projection images. Maximum intensity projections of the heart of *Tg(myl7:ERK-KTR-Clover-p2a-H2B-mScarlet)* larvae at 72 hpf. **(B)** Larvae treated with the Erk inhibitor exhibit an increased number of CMs with inactive Erk signaling (arrows) compared with larvae treated with DMSO **(A)**. **(C)** Quantification of ventricular CMs with nuclear Clover. All images are ventral views, anterior to the top. V, ventricle. Data in graphs expressed as mean ± SEM.

**Figure 6-figure supplement 2.**
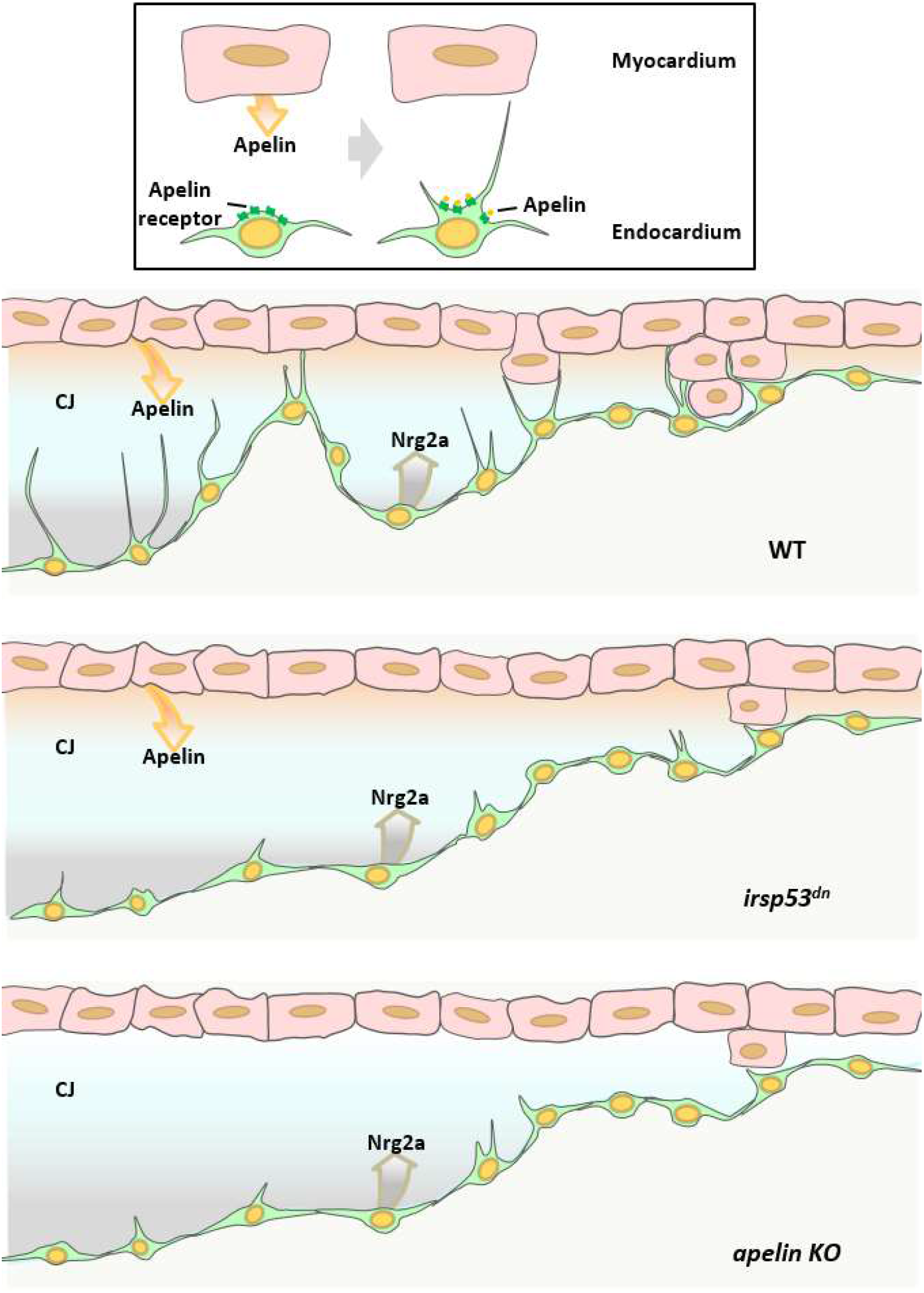
Schematic model. Schematic model depicts that manipulating the formation of endocardial protrusions results in cardiac trabeculation defects via mediating the function of Nrg/ErbB signaling.

## Notes

### Competing Interest Statement

The authors have declared no competing interest.

